# Neuronal Avalanches in Naturalistic Speech and Music Listening

**DOI:** 10.1101/2023.12.15.571888

**Authors:** Matteo Neri, Claudio Runfola, Noemie te Rietmolen, Pierpaolo Sorrentino, Daniele Schon, Benjamin Morillon, Giovanni Rabuffo

## Abstract

Neuronal avalanches are cascade-like events ubiquitously observed across imaging modalities and scales. Aperiodic timing and topographic distribution of these events have been related to the systemic physiology of brain states. However, it is still unknown whether neuronal avalanches are correlates of cognition, or purely reflect physiological properties. In this work, we investigate this question by analyzing intracranial recordings of epileptic participants during rest and passive listening of naturalistic speech and music stimuli. During speech or music listening, but not rest, participants’ brains “tick” together, as the timing of neuronal avalanches is stimulus-driven and hence correlated across participants. Auditory regions are strongly participating in coordinated neuronal avalanches, but also associative regions, indicating both the specificity and distributivity of cognitive processing. The subnetworks where such processing takes place during speech and music largely overlap, especially in auditory regions, but also diverge in associative cortical sites. Finally, differential pathways of avalanche propagation across auditory and non-auditory regions differentiate brain network dynamics during speech, music and rest. Overall, these results highlight the potential of neuronal avalanches as a neural index of cognition.

**Author’s summary:** Neuronal avalanches consist of collective network events propagating across the brain in short-lived and aperiodic instances. These salient events have garnered a great interest for studying the physics of cortical dynamics, and bear potential for studying brain data also in purely neuroscientific contexts. In this work we investigated neuronal avalanches to index cognition, analyzing an intracranial stereo electroencephalography (iEEG) dataset during speech, music listening and resting state in epileptic patients. We show that neuronal avalanches are consistently driven by music and speech stimuli: avalanches co-occur in participants listening to the same auditory stimulus; avalanche topography differs from resting state, presenting partial similarities during speech and music; avalanche propagation changes during speech, music, and rest conditions, especially along the pathways between auditory and non auditory regions. Our work underlines the distributed nature of auditory stimulus processing, supporting neuronal avalanches as a valuable and computationally advantageous framework for the study of cognition in humans.

## Introduction

The brain is a complex system operating across multiple spatial and temporal scales. In order to decode brain functions, researchers analyze brain data across imaging modalities searching for regularities, such as patterned processes that might correlate with specific behaviors or cognitive abilities (1,2). Recurrent patterns can be observed, for instance, in the spectral analysis of brain oscillations (3,4) or in the investigation of functional connectivity (5). One ubiquitous finding, observed across imaging modalities and spatial and temporal scales, is the spontaneous occurrence of cascade-like events recurring aperiodically in neural activity (6–10). Historically, the interest in the study of these events originated from the observation that their statistics (e.g., the seemingly scale-free distribution of event duration and size) drew parallels with natural phenomena like sandpile avalanches and forest fires (11). For this reason, they were termed *neuronal avalanches*. Neuronal avalanches have been hypothesized to be the manifestation of a critical state of neural dynamics, corresponding to neural activity occupying a sweet spot at the edge of two different regimes of organization (e.g., synchronous and asynchronous (12,13)). Brain criticality was proposed as a general organizing principle for the brain (14) and has been hypothesized to play a crucial role in information processing (15), adaptive behavior (7,16), and robustness of brain networks (17,18). While the brain criticality hypothesis remains debated (19), evidence shows (20) that neuronal avalanches are a valid marker of physiological brain states beyond the criticality assumption. For instance, neuronal avalanche features differ across health and pathology (21–25) or across resting wakefulness and sleep states (8,26). Most studies in humans have analyzed avalanches in task-free conditions. An open question hence remains about whether neuronal avalanches index cognitive functions or purely reflect physiological states.

Recent literature has argued that applying analytical frameworks used in resting-state studies to cognitive task datasets could elucidate how cognition emerges from neuronal activities (27). Within this context, experiments based on naturalistic stimuli offer a convenient trade-off between overly specific task paradigms and non-specific resting-state studies (28–31). The use of ecologically valid stimuli—both in terms of content and duration—makes it possible to study changes in neuronal avalanche properties in continuous brain activity induced by specific natural cognitive states (29,32,33). This allows investigating the hypothesis that avalanches are a neurophysiological correlate of cognitive processes and, as such, that their properties are modulated according to the stimulus at hand. To test this, we analyzed neuronal avalanches recorded in 19 pharmaco-resistant epileptic participants using intracranial electroencephalography (iEEG) recordings, with channels distributed across the entire brain (Fig.1.A). In particular, we focussed on high-frequency activity (HFa, 80-120 Hz), because it is an effective index of local neuronal processing and is highly correlated with local neuronal spiking activity (34). Such recordings provide anatomically precise information about the functionally-selective engagement of neuronal populations at the millimeter scale as well as about their temporal dynamics at the millisecond scale. This setup is essential for accurately portraying the neurophysiological underpinning of a specific cognitive process (35,36). To establish ecologically-relevant stimulation conditions, participants listened to long and natural audio recordings of either speech or music and also underwent a control resting-state (rest) session (∼10 minutes per condition). Given our hypothesis, we expect that the neuronal avalanche timing and propagation across the brain is affected by the presence and the nature of the stimuli (16). For each condition, we analyzed the temporal and topographic properties of neuronal avalanches, showing that these salient events synchronize across participants listening to the same auditory stimulus, revealing common neural patterns associated with processing of temporally structured musical and linguistic patterns. We show that the co-occurrence of avalanches across participants indexes cognitive processes over and beyond auditory perception, retrieving significant inter-subject correlations without considering the auditory regions and with sparse channel selection. Then, to gain further insights into speech and music processing we tracked the propagation of avalanches spatially, and studied which regions participate in coordinated neuronal avalanches, and how music and speech alter the spread of avalanches propagation across the brain. Focusing on neuronal avalanches allowed us to unveil, with low computational cost, new insights about brain function (37). Our analysis supports the idea that neuronal avalanches are network-level correlates of cognitive processes capturing both distributed and specific functional properties.

**Figure 1.**
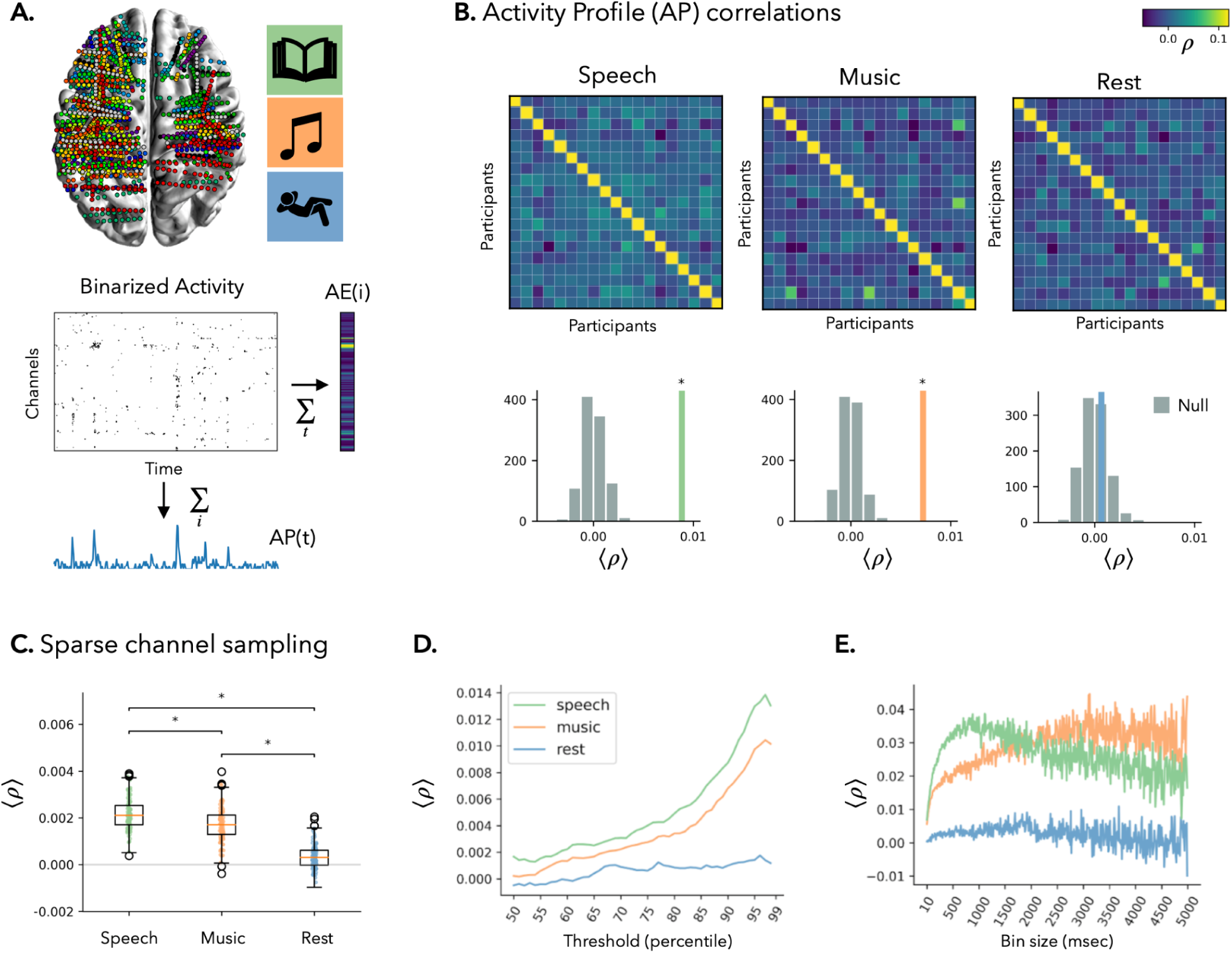
**A) (top left)** Anatomical localization of the iEEG electrodes for each participant (N = 19); different colors correspond to different participants. **(top right)** The three natural conditions under study: speech, music and rest. **(bottom)** Binarized HFa over time, for the different iEEG channels of a representative participant. The Activity Engagement (AE) and the Activity Profile (AP) are obtained by summing the binarized activity matrix across time and across channels, respectively. **B) (top)** Inter-subject correlation matrix for the three conditions. Each point of the matrix represents the correlation value between the APs of two participants. **(bottom)** Mean of the inter-subject correlation (colored line) compared to a null model distribution (see Methods; *P<0.001). **C)** Distribution of mean inter-subject correlation obtained by randomly selecting one channel per anatomical region (1000 iterations) in all the participants. The distributions for speech, music, and rest are compared using the Wilcoxon signed-rank test (*P<0.001). **(D-E)** Average inter-subject correlation computed by binarizing the data with **(D)** different thresholds and **(E)** bin sizes (see Methods).

## Results

### Neuronal avalanches support inter-subject correlations during speech and music

Neuronal avalanches were estimated by binarizing the z-scored HFa and selectively focusing on salient—above threshold (99 percentile)—events in the neuronal dynamics (Fig.1.A; see Methods). This approach captures higher-order neuronal dynamics while being a sparse code and hence being computationally efficient. Neuronal avalanches are then defined as consecutive above-threshold activations and correspond to collective network events occurring across subsets of recording channels. As a convention, an avalanche starts when at least one channel is ‘active’, and ends when no channels are above the threshold.

We first analyzed the timing of neuronal avalanches, defining their activity profile (AP) as the sum of the binarized data across channels (Fig.1.A), which measures the number of channels participating in a neuronal avalanche at each time-point. We computed the inter-subject correlation of the APs over time, for each condition (Fig.1.B; see Methods). We observed a significant inter-subject correlation between the APs of different participants during both speech and music listening, but not during rest (permutation tests: speech: p < 0.001; music: p < 0.001; rest: p = 0.29; Fig.1.B). The APs extracted from only the auditory cortex were significantly correlated across participants listening to the same acoustic stimulus, indicative of a stimulus-driven response (speech: p < 0.001; music: p < 0.001; rest: p = 0.062; Supp. Fig.S1.A). Moreover, when considering only the APs of a distributed network of non-auditory associative cortical regions, a significant inter-subject correlation was observed for speech, but not for music nor for rest (speech: p = 0.007; music: p = 0.09; rest: p = 0.41; Supp. Fig.S1.B). In order to exclude that the inter-subject correlation was a byproduct of synchronous activations of spatially contiguous channels, we evaluated the inter-subject correlation for random picks of recording channels belonging to different anatomical regions (hence being spatially distant). Also in this case, a non-trivial correlation was observed during speech and music (but not rest) conditions (speech: p < 0.001; music: p = 0.003; rest: p = 0.261; Fig.1.C; see Supp. Fig.S1.C for assignment of channels to anatomical regions). Our results highlight that speech and music processing are distributed processes, involving networks of brain regions activating together in response to external stimuli (38).

The binarization step requires fixing two main parameters: the threshold above which a channel is considered active, and a bin size that is used for temporally coarse-graining the data (see Methods). Thus, in order to check for the stability of our results we computed the inter-subject correlation for different values of threshold and bin size parameters. We observed that the thresholds at which the inter-subject correlation is maximal for both speech and music — and mostly different from rest— are above 95 percentile (Fig.1.D). This shows that focusing on rare salient events (i.e., using a high threshold when binarizing), while computationally more efficient, also selects features that map more closely on cognitive processes in the brain.

The binning process defines a temporal coarse-graining of our data, obtained by dividing each channel’s binary activity in time bins of size *m* and assigning 1 to bins when at least one activation was observed, and 0 if the channel was never active during the *m* time steps. The binning step allows us to look at the data with different time resolutions, and it is often used in the literature e.g., to search for evidence of criticality in brain data (39), avoiding random noise effects in the scale-free distribution of neuronal avalanches size and duration. Here, we observe a peak in the inter-subject correlation for speech at bin size *m ⋍ 100,* corresponding to ∼1 second (Fig.1.E, green). For music, the inter-subject correlation reaches its maximal values at longer temporal scales around ∼3 seconds (Fig.1.E, orange). This suggests that these two conditions may be characterized by different time scales of neuronal processing. As expected, the mean inter-subject correlation remains at baseline during rest, regardless of the bin size (Fig.1.E, blue). In the following analyses, we fixed *m=2* (and threshold = 99 percentile). This way, our results are not biased by the characteristic timescales of speech and music processing, while maintaining a high temporal resolution.

### Neuronal avalanches are distributed during speech and music processing

We performed the topographic analysis of neuronal avalanches during speech, music, and rest, by analyzing the Activity Engagement (AE), a metric obtained by summing the binarized data over time (Fig.1.A), and quantifying the number of times each region is recruited by an avalanche event (a similar measure is also adopted in (16)). We hypothesized that the AE would differ across conditions. To test this, for each participant and for each channel *i*, we compared the AE(*i*) across pairs of conditions (i.e., speech versus rest, music versus rest, and speech versus music). Statistical significance was assessed by comparing the observed AE(*i*) against a corresponding theoretical null-distribution, computed under the null hypothesis that no difference in AE(*i*) would occur across any pair of conditions A and B (see Methods). We define “high-AE channels” those channels where the AEs is significantly higher when evaluated against the null-distribution (with p < 0.05, Bonferroni corrected for the number of channels; Fig.S2.A).

This analysis reveals that channels engaged during speech and music processing are spatially distributed and mostly non-overlapping between conditions (Fig.2A). In addition, several complementary channels disengage compared to the resting state baseline (Fig.S2.B). Importantly, the analysis of activity engagement (AE) is complementary to the analysis of HFa, as these metrics are poorly correlated (Fig.S3), and HFa has been shown to be spatially overlapping between the speech and music conditions (3).

**Figure 2.**
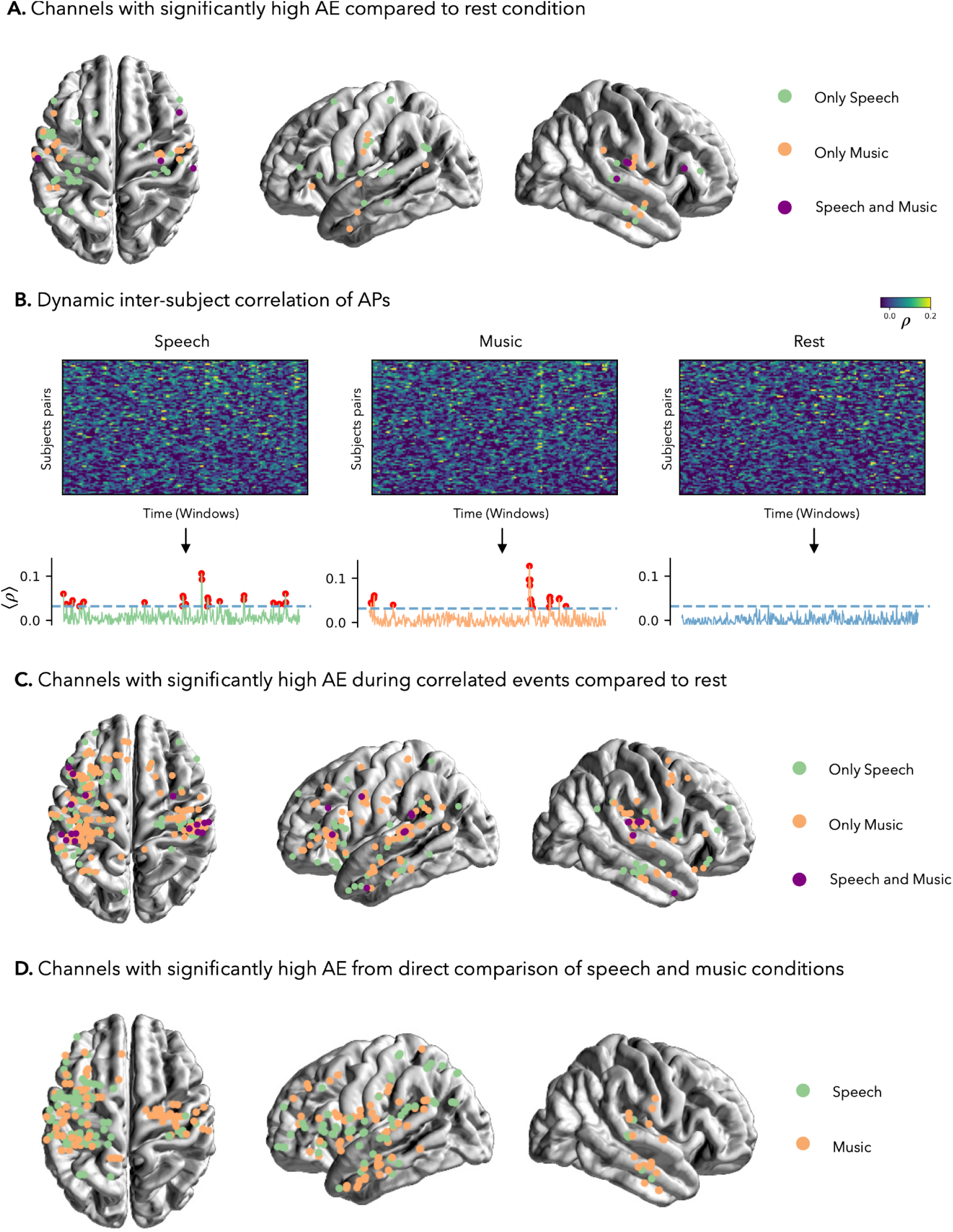
**A)** Channels with significant activity engagement (AE, i.e., the sum over time of the binarized activations) compared to rest. Channels in green (/orange) mark significant AE during speech (/music) compared to rest, and channels in purple mark significant AE during both speech and music when independently compared to rest. **B) (top)** Dynamic inter-subject correlation of APs plotted per time window (1 sec) and for each pair of participants. **(bottom)** Evolution over time of the mean inter-subject correlation during speech (green), music (orange) and rest (blue). The red dots correspond to the windows of time in which the inter-subject correlation is higher than the maximum observed during rest (dashed blue line). **C)** AE computed during moments of high inter-subject correlation (red dots in panel A). The same conventions as in panel A are adopted. **D)** Direct comparison of AE between the speech and music conditions (see Methods). **A-D)** p <0.05, Bonferroni corrected for the number of channels.

### Neuronal avalanches topographies during strong inter-subject correlation

To understand whether inter-subject correlations of APs are driven by specific coordinated events, we correlated the APs in a time-resolved way, using a sliding window approach with 1 second window size (see Methods). The mean inter-subject correlation value fluctuates over time (Fig.2.B), suggesting that there are specific moments in which neuronal avalanches of different participants listening to the same stimulus co-occur. In several time windows during speech and music listening, the inter-subject correlations indeed display high values, above the maximum observed during rest (Fig.2.B, bottom). The topographic analysis of neuronal avalanches during time windows of high inter-subject correlation (i.e., > rest; Fig.2.C) reveals more “high-AE channels” during naturalistic speech or music listening as compared to rest conditions (see Table.S1). Significantly higher engagements are observed in channels distributed over the entire cortex. These channels are partially overlapping between speech and music, mostly within the auditory cortex, but also in associative temporal and frontal regions. These results indicate that the inter-subject correlation of APs during speech and music processing is driven by whole-brain network phenomena, not limited to local (auditory) processing.

### Partial overlap of neuronal avalanches during speech and music conditions

Finally, we performed the topographic analysis of AE by directly comparing speech versus music conditions, which reveals a different localization of high-AE channels during speech and music processing (Fig.2.D). While our analysis highlights differences in the networks engaged during speech and music listening, the percentage of channels for which the AE is significantly different when comparing speech versus music is low (6.3% of the channels for the aggregated population; see Table.S1). This result is even more marked at the individual level, as most participants display a remarkably low percentage of channels with significant high-AE (mean-percentage across participants = 2.83% , std=4.69). This result supports previous evidence that speech and music processing mostly rely on shared resources (3). Similar low percentages are observed when comparing speech versus rest, and music versus rest (Table.S1), which suggests specific localized alterations of brain activity during speech/music processing compared to rest condition.

### Neuronal avalanche propagation pathways change between conditions

In order to further investigate the speech, music and rest conditions, we analyzed the spatiotemporal propagation of neuronal avalanches through the brain. We computed the avalanche transition matrices (ATMs) (22,40), which measure the probability that the activation of channel *i* results in the activation of channel *j* at the successive time step (Fig.3.A; see Methods). Hence, for each participant, the ATMs estimate the characteristic pathways through which neuronal avalanches propagate. We computed the ATMs for each condition (example participant in Fig.3.B) and compared them by correlating their respective (vectorized) ATMs (Fig.3.C). The ATMs of rest (r) and music (m) are significantly more correlated than the ATMs of speech (s) and rest, or speech and music (Wilcoxon test: p = 0.003 for (s,r) versus (r,m); p = 0.86 for (s,r) versus (s,m); p = 0.006 for (r,m) versus (s,m)). This suggests that avalanche propagation during music listening and rest is more similar than during speech listening.

**Figure 3:**
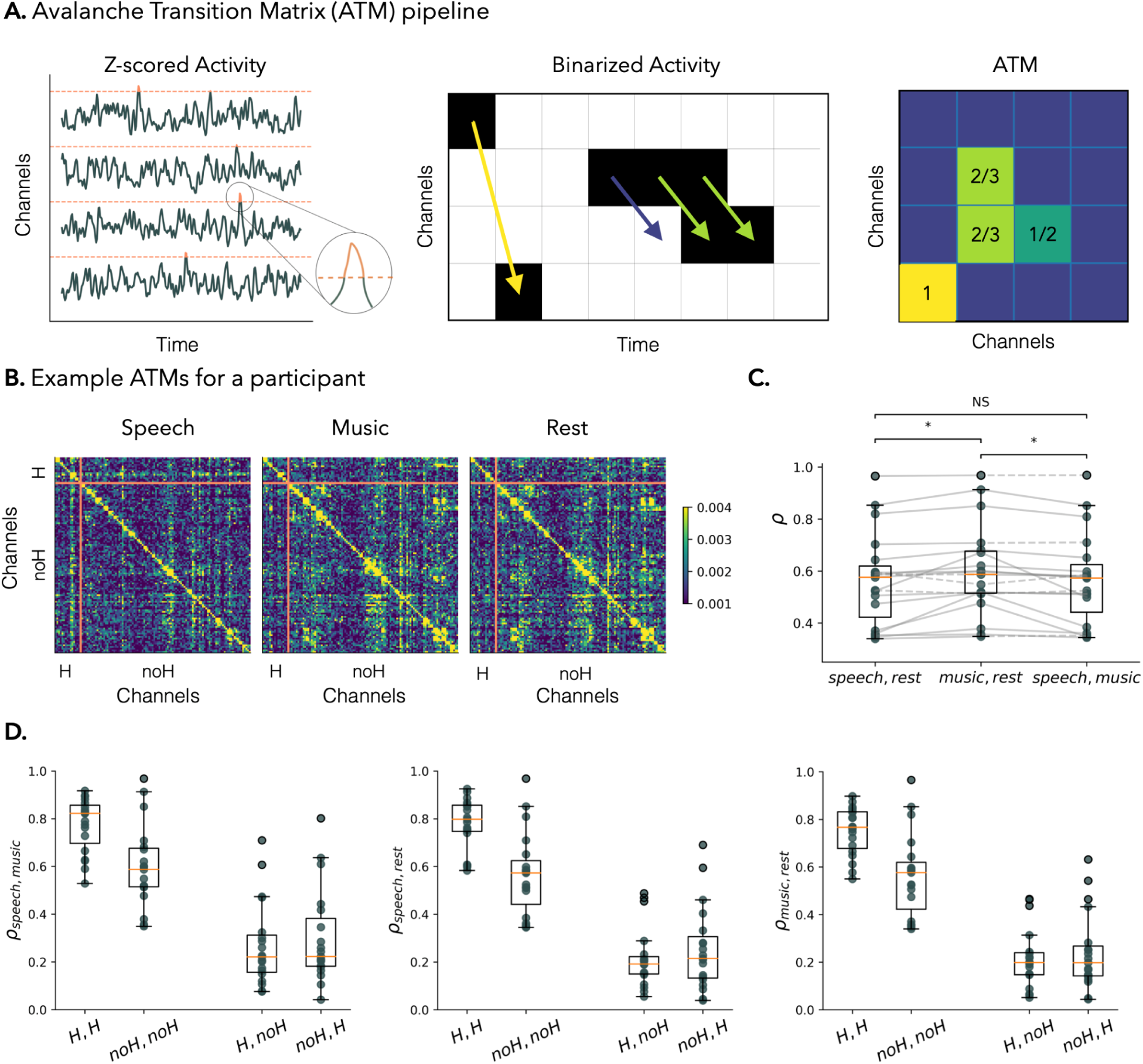
**A)** Avalanche transition matrices (ATMs) pipeline. **(left)** example of binarized activity extracted from the HFa. **(middle)** For each pair of channels i,j of a participant, the transition of activity between channel i at time t and channel j at time t+1 is estimated. **(right)** Each element in position i,j of the ATM represents the probability that channel j records a salient activation at time t+1 given that channel i recorded a salient activation at time t. **B)** ATMs of one participant for the three conditions. The orange lines separate the four subsets of ATMs’ elements investigated (H-H, noH-noH, noH-H, H-noH). **C)** Distribution of Pearson’s correlation values (ρ) across participants, computed between ATMs of pairs of conditions. Solid/dashed gray lines: participants showing the group-average/reversed effect (Wilcoxon test: *p < 0.01; NS non-significant). **D)** Distribution of correlation values across participants, computed between ATMs of pairs of conditions. ATMs were estimated within (H-H, noH-noH) and between (H-noH; noH-H) areas. The boxplots in panels C-D represent the median and the interquartile range. H: Auditory (Heschl’s gyrus); no-H: non-auditory channels.

To disentangle the contribution of local auditory (intra-areal) and distributed (inter-areal) processes we separated auditory (Heschl’s gyrus, H) and non-auditory (noH) channels, and estimated the ATMs within (H-H, noH-noH) and between (H-noH; noH-H) them. Note that, as ATMs estimate directional transitions, the H-noH ATM refers to avalanches projecting from auditory to non-auditory regions, while the noH-H ATM to neuronal avalanches incoming to the auditory cortex. Next, we analyzed the correlations between the ATMs by focusing on the H-H, noH-noH, H-noH, and noH-H blocks separately.

The correlations between the ATMs in H-H and noH-noH are higher than those considering H-noH and noH-H blocks, consistently across all pairs of conditions (Fig.3.D; comparing in each panel the first two distributions to the last two; paired non-parametric Wilcoxon signed-ranks tests: all p < 0.001; all z-statistic >3.7). This indicates that the functional connections that are changing the most between conditions are the ones between auditory and non-auditory regions (either projecting from or going into H-channels), while the connections between channels localized both within (H-H) or outside (noH-noH) the auditory cortex are more stable across conditions. Moreover, the high correlation values of ATMs in H-H across all pairs of conditions suggest that the transition matrix does not vary much within the auditory cortex across conditions. Correlations in H-H are higher than those in noH-noH; all p < 0.002; all z-statistic > 3.1.

Finally, the correlations between the ATMs in H-noH and noH-H are significantly different only when comparing speech and music (Fig.3.D, left panel; p = 0.036), but they are not statistically different when comparing speech to rest or music to rest (Fig.3.D, central and right panels; p < 0.08). In all cases, the ATMs correlations in the H-noH and noH-H blocks are rather low (Pearson’s ρ ∼ 0.2). These results indicate that the (feedforward and feedback) propagation of neuronal avalanches between auditory and associative regions best indexes the specificity of cognitive processes induced by different conditions.

## Discussion

Naturalistic stimuli such as movies, virtual reality, speech, or music listening, create an ecologically-relevant set up to study the emergence of neuronal processes and the mechanisms supporting brain functioning. In the present work, we investigated two major auditory cognitive domains, namely speech and music processing. As both domains are universally and uniquely human, scientists, since Rousseau and Darwin, have debated whether speech and music share common or different origins. In neuroscience, this debate can be addressed by assessing how these processes are dynamically mapped into brain activities.

Brain mapping requires processing and analyzing brain signals, for which there exists no unique approach. In fact, as a complex system, the brain can be effectively studied according to different perspectives. A canonical perspective to interpret brain data is in terms of brain rhythms or oscillations (41,42). Temporal analysis and spectral decomposition of “analog” signals allow isolating frequencies of interest with a putative role in a specific brain function (43). A complementary perspective to study information processing in the brain is offered by the analysis of salient events, which generally occur in aperiodic bursts (44). In fact, from the early cybernetic interpretation of the brain as a “digital” machine (45) to the more recent applications of information theory tools (46), it was shown that a great deal of information can be extracted by binarizing the signals. In this work, we demonstrated this principle by analyzing an iEEG dataset during naturalistic acoustic stimuli preserving only a small percentage of data points corresponding to neuronal avalanches. This approach effectively retrieved relevant information about brain processing during speech and music listening, including intersubject correlations (Fig.1), and characteristic topographies of activations (Fig.2-3).

Binarizing signals presents a compelling computational advantage, demanding significantly lower computational resources compared to the analysis of temporal and spectral signal properties. Notably, the binarization process employed in this work retains only approximately 2% of the data points. Despite this substantial reduction in the volume of data considered, the predictive power of binarized activity remains robust. In fact, decreasing the amount of data can enhance the quality of our results, as illustrated in Fig.1.D, where higher binarization threshold led to stronger inter-subject correlations. This underscores the efficiency and effectiveness of signal binarization in optimizing computational resources without compromising analytical insights.

Even though in each participant the channels were placed in different locations, we found significant inter-subject correlations of the APs, suggesting that the processing of speech and music in the human brain do not involve only some specialized regions, but rather a degenerate computation (47,48) involving different brain areas sharing similar temporal dynamics. This result is further strengthened by our analysis performed on a sparse channel selection, where using only one channel per anatomical region preserved significant inter-subject correlations (Fig.1.C).

Our experiment, conducted in ecologically valid conditions, allowed participants to naturally process speech and music across various levels, from phonemes to sentences, and from single notes to musical phrases. During speech listening we observed significant inter-subject correlation also after excluding the channels in auditory regions (Fig.S1). This might reflect integrative “high-level” stimulus-driven cognitive processes during speech listening. While this was not observed during music, we expect that a temporal coarse graining would reveal a similar feature also during music listening, in virtue of the better performances shown by large binning values during music listening (Fig.1.E). Alternatively, the divergence could stem from individualized processing of music phrases as compared to speech sentences, where the linguistic/semantic processing during storytelling temporally aligns across subjects, while personal experiences differ across subjects listening to music.

These results confirm previous work that describe speech and music processing as distributed across the whole-brain (49,50). However, it is still unclear whether these processes are related to language/music specific regions or if, rather, they rely on shared networks (51). Previous work on the same dataset analyzed in this manuscript showed that, in terms of both HFa and other frequency bands, the spatial specializations for speech and music are in fact alike: they do not recruit selective cortical regions but mostly rely on shared resources (3). Our analysis of neuronal avalanches topographies (using the rest condition as a baseline) partially confirms these results, as several channels with high-AE were shared between speech and music processing (Fig.2.A,C, purple channels). This was particularly evident in auditory areas, though not exclusively. Away from these areas, music and speech were characterized by high-AE in different subnetworks (Fig.2.A,C). This confirms that these two conditions rely on partially overlapping whole brain networks hinging across different brain areas. A direct comparison of the speech and music conditions confirms the differences between speech and music processing (Fig.2.D), but only in a small percentage (less than 3%) of the channels.

Finally, examining the propagation of neuronal avalanches (Fig.3), we discovered that connections between auditory and non-auditory regions exhibit the least correlation across different conditions. This implies that the most dynamic changes between rest, speech, and music occur along the feedforward and feedback pathways. Additionally, functional connections outside the auditory regions demonstrate significantly lower correlation across conditions compared to those within. This shows that speech and music listening robustly modulate the information flow among frontal and temporal cortical regions, and less so in the auditory cortex itself.

In this work, we focused on (the binarization of) HFa signals, which are generally used to provide insights into the neural bases of cognition (52). In general, the analysis of binarized signals offers a complementary view of brain data, which is only weakly related to more diffuse measures of brain oscillations (e.g., signal power; Fig.S3). Typically, above-threshold activities are interpreted as higher-order dynamical properties. In appendix A and Fig.S4 we further show that the AP and AE measures introduced in this work, and obtained from the binarized activity, can be associated with the characteristic decay time of signals’ autocorrelation,

In appendix A and Fig.S4 we further show that the AP and AE measures introduced in this work, and obtained from the binarized activity, can be associated with the characteristic decay time of the signals autocorrelation, a measure that in the “analog” signal processing framework has been used to measure the presence of characteristic timescales in brain activity. In this context, the “digital” framework of the AP and TP could offer a time-resolved and computationally advantageous proxy of timescales dynamics. Besides computational efficiency, binarization allows the deployment of a series of conceptual tools such as information theory (46), the study of network dynamics (18), as well as paradigms such as brain criticality (53,54). In relation to the latter, we highlight that, in standard neuronal avalanches analyses aimed at testing the presence of a critical state in the brain, the bin size value chosen for the data binarization plays an important role. In fact, the tuning of the bin size value is important to guarantee the observation of power-law distributed neuronal avalanches (39). Analyzing the intersubject correlations for different bin size values (Fig.1.E), we showed that speech and music processing are characterized by different timescales, with optimal performances for different bin size values. This suggests that, depending on the problem at hand, a convenient definition of the bin size can also be determined by functional performances, rather than by the search for any particular statistics of neuronal avalanches.

In conclusion, our study provides compelling evidence that neuronal avalanches offer an effective means to characterize neuronal processing in naturalistic conditions. This provides a valuable complement to oscillation-based frameworks, thereby enriching the ongoing discourse on the neuronal underpinnings of speech and music processing.

## Materials and Methods

### Participants

19 participants (10 females, mean age 30 y, range 8-54 y) with pharmacoresistant epilepsy participated in the study. All patients were French native speakers. Neuropsychological assessments, carried out before stereotactic intracranial EEG (iEEG) recordings, indicated that all participants had intact language functions and met the criteria for normal hearing. In none of them were the auditory areas identified as part of their epileptogenic zone, as per the rating of an experienced epileptologist. Recordings took place at the Hôpital de La Timone (Marseille, France). Patients provided informed consent prior to the experimental session, and the experimental protocol was approved by the Institutional Review board of the French Institute of Health (IRB00003888).

### Data acquisition

The iEEG signal was recorded using depth electrodes shafts of 0.8 mm diameter containing 10 to 15 electrode contacts (Dixi Medical or Alcis, Besançon, France). The contacts were 2 mm long and were separated from each other by 1.5 mm. The locations of the electrode implantations were determined solely on clinical grounds. Participants were included in the study if their implantation map covered at least partially the Heschl’s gyrus (left or right). The cohort consists of 13 unilateral implantations (10 left, 3 right) and 6 bilateral implantations, yielding a total of 271 electrodes and 3371 contacts (see Fig.1.A and S1.C for electrodes localization).

The patients were recorded either in an insulated Faraday cage or in the bedroom. In the Faraday cage, participants sat comfortably in a chair, the room was sound attenuated, and data were recorded using a 256-channels amplifier (Brain Products), sampled at 1kHz and high-pass filtered at 0.016 Hz. In the bedroom, data were recorded using a 256-channels Natus amplifier (Deltamed system), sampled at 512 Hz and high-pass filtered at 0.16 Hz.

### Experimental design

The participants passively listened to 10 minutes of storytelling (576,7 secs, La sorcière de la rue Mouffetard, ref Gripari P. 2004) and 10 minutes of music (580.36 secs, Reflejos del Sur, ref Oneness, 2006) embedded in 3 resting-state sessions (rest; each lasting 3.3 minutes). The order of the speech and music conditions was balanced across participants. In all our analyses, we concatenated the 3 resting-state sessions into a single 10-minute-long rest condition. In the Faraday cage, a sound Blaster X-Fi Xtreme Audio, an amplifier Yamaha P2040, and Yamaha loudspeakers (NS 10M) were used for sound presentation. In the bedroom, stimuli were presented using a Sennheiser HD 25 headphone set. Sound stimuli were presented with a 44.1 kHz sample rate and 16 bits resolution. Speech and music excerpts were presented at ∼75 dBA.

### Preprocessing

Contact data were converted offline to virtual channels using a bipolar montage approach (closest-neighbor contact reference) to increase spatial resolution and reduce passive volume diffusion from neighboring areas (55). To precisely localize the channels, a procedure similar to the one used in the iELVis toolbox was applied (56). Anatomical localization of the channels were further labeled with the Brainnetome Atlas (Fig.S1.C; (57)).

Bipolar channels outside the brain were removed from the data (∼3 %). The remaining data were bandpass filtered between 0.1 Hz and 250 Hz and, when necessary, a notch filter was additionally applied at 50 Hz and harmonics up to 200 Hz, to remove power line artifacts. Finally, the data were downsampled to 500 Hz.

### High-frequency activity (HFa)

The amplitude of the high-frequency activity (HFa, 80-120 Hz) was obtained by computing —via the Hilbert transform— the analytic amplitude of four 10-Hz-wide sub-bands spanning from 80 to 120 Hz. Each sub-band was standardized by dividing it by its mean and, finally, all sub-bands were averaged together (58); (59). To further reduce the computational cost, the data were downsampled to 100 Hz.

### Artifact rejection

Channels with a variance greater than 2*IQR (interquartile range, i.e., a non-parametric estimate of the standard deviation), were tagged as artifacted channels. Then, for the remaining channels, time-segments with activity greater than 5 standard deviations were considered to be artifacted and replaced with the mean of the signal. This prevented destroying the temporal structure of the data, which is necessary to evaluate the inter-subject correlations.

### Activity profile (AP)

The activity profile (AP) reflects the timing of neuronal avalanches, and is estimated as the sum of the binarized HFa across channels (Fig.1.A). It measures the number of channels participating in a neuronal avalanche at each time point. To this end, HFa signals *X_i_* (*t*) were first z-scored across time, as, 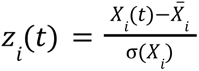; where *X̄* and σ(*X*) denote the mean and variance of the signals, respectively. The binarization was then performed by dividing the data in bins of *m* time steps. If a bin contains at least one time-point in which the activity in absolute value is higher than the fixed threshold, the bin is assigned value 1, and 0 otherwise. The threshold was fixed as the 99th percentile of the (absolute value of) z-scored resting-state activity, and the same threshold was used for all conditions, i.e., speech, music and rest. Fig.1.D shows that such a high threshold value guarantees optimal inter-subject correlation results, in line with the idea that brain dynamics is largely driven by rare activation events. Unless differently specified, we fixed *m*=2 to guarantee high temporal resolution while reducing noise effects. Notice that the tuning of the bin size *m* can play an important role in detecting critical-like avalanches (39). To check for the stability of our results, in Fig.1.E we computed the inter-subject correlation using different *m*, which provides indirect insights about the temporal scales of speech and music processing.

### AP correlations

The inter-subject correlation of the APs over time was obtained by computing the correlation between the APs of each pair of participants and averaging across all the pairs. To test whether inter-subject correlations are statistically significant, we generated null models by randomly shifting the AP of each participant before computing the inter-subject correlation. We iterated this process 1000 times and we compared the null-distributions of the inter-subject correlations with the observed inter-subject correlations. To establish significance, we computed p values as the proportion of iterations in which the correlation in the surrogate data was higher than the observed one. To test that inter-subject correlations are not exclusively driven by local interactions, i.e., within channels situated in the same cortical regions, we sampled randomly one channel per atlas-based region in all the participants. We calculated the correlation between the APs, computed using only the sampled recording sites. We iterated this process 1000 times and we established the significance of our results computing p values as the fraction of samples (in total 1000) in which a positive inter-subject correlation was measured. The distribution of the 1000 results are compared across conditions using the Wilcoxon signed-rank test (Fig.1.C).

### Dynamic inter-subject correlation

To investigate the temporal evolution of the inter-subject correlation we correlated the APs in a time-resolved way, using a sliding window approach. That is, we looked at the evolution of inter-subject correlation between APs across time windows of 50 time points (1 seconds). To maintain a good temporal resolution we used a sliding step equal to the temporal resolution of the binarized time series, i.e.20 milliseconds. This way, we obtained, for each condition, a time series for the inter-subject correlation.

### Activity Engagement (AE)

In order to perform a topographic analysis of neuronal avalanches across speech, music, and rest conditions we defined the activity engagement (AE) as the sum over time of the binarized signals (Fig.1.A). Then, for each brain region *i*, we assessed the significance of the observed AE(*i*) by pairwise comparisons of the speech, music, and rest conditions .

Given any two conditions A and B, we defined a null-distribution of AE_*Null*_(*i*) based on the null-hypothesis that AE_A_(*i*) and AE_B_(*i*) are not different from each other. According to this, starting from the *i*-th binary signal of time length *n* in conditions A and B, we defined a new binary sequence by randomly selecting *n* time bins from the joint A and B signals. The sum of this new sequence defines a “null” AE for the *i*-th channel. This process can be repeated a large number of times to build a null-distribution of AE_*Null*_(*i*) values. For a large numbers of iterations this distribution approaches a hypergeometric distribution with mean *k*=(AE_A_(*i*)+AE_B_(*i*))/2 and variance *k*(*n* − *k*)/(2*n* − 1). This allowed us to efficiently perform statistical testing by comparing AE in two conditions against the expected theoretical distribution, avoiding computationally expensive iterations. We defined as “high-AE channels” those channels with significantly higher AE as compared to the theoretical null-distribution (*P* < 0. 05, Bonferroni correction for the number of channels).

We also utilized this approach to compare the AEs for speech and music computed exclusively over the time steps belonging to the windows of times in which the intersubject correlation for speech and music was higher than the maximum observed in resting state (Fig.2.B-C).

### Avalanches transition matrix (ATM)

We studied the propagation patterns of avalanches computing the ATM, from the recorded activity in a similar way as in (40). Following the estimation of the APs described above, we focused on single neuronal avalanches, defined as sequences of continuous activity (i.e., when at least two consecutive time steps of the AP are non-zero) starting after and ending before a zero-value bin of the AP. We first constructed an avalanche-specific transition probability wherein the (*i,j)* entry represents the probability that channel *j* is above threshold (i.e., it is 1 in the binarized data), at time *t+1* given that channel *i* was above threshold at time *t.* The participant- and condition-specific ATM is then built as the mean of the previously computed matrices across avalanches. A scheme of the ATM computation process is illustrated in Fig.4A.

ATMs were estimated for each condition at the whole brain level, but also analyzed by separating auditory (Heschl’s gyrus, H) and non-auditory (nonH) channels to estimate ATMs within (H-H, nonH-nonH) and between (H-nonH; nonH-H) them. Auditory channels corresponded to all the channels within the electrode that was traversing the Heschl’s gyrus, as defined visually by the neurosurgeon during the implantation procedure. The similarity of the ATMs between conditions was estimated as the absolute value of the Pearson’s correlation between the vectorized (excluding the main diagonal) ATMs. The stability of the distributions of the correlations was tested using a bootstrap approach. Statistical significance of the difference of correlation between pairs of conditions was evaluated via the Wilcoxon test.

## Acknowledgments

This work was supported by the European Union (ERC, SPEEDY, ERC-CoG-101043344), ANR-16-CONV-0002205 (ILCB) and the Excellence Initiative of Aix-Marseille, University (A*MIDEX). Matteo Neri would like to thank Giovanni Petri, Andrea Brovelli, Etienne Combrisson and Bruno Giordano for the discussions we had.

## Supplementary Material

**Supplementary Figure S1.**
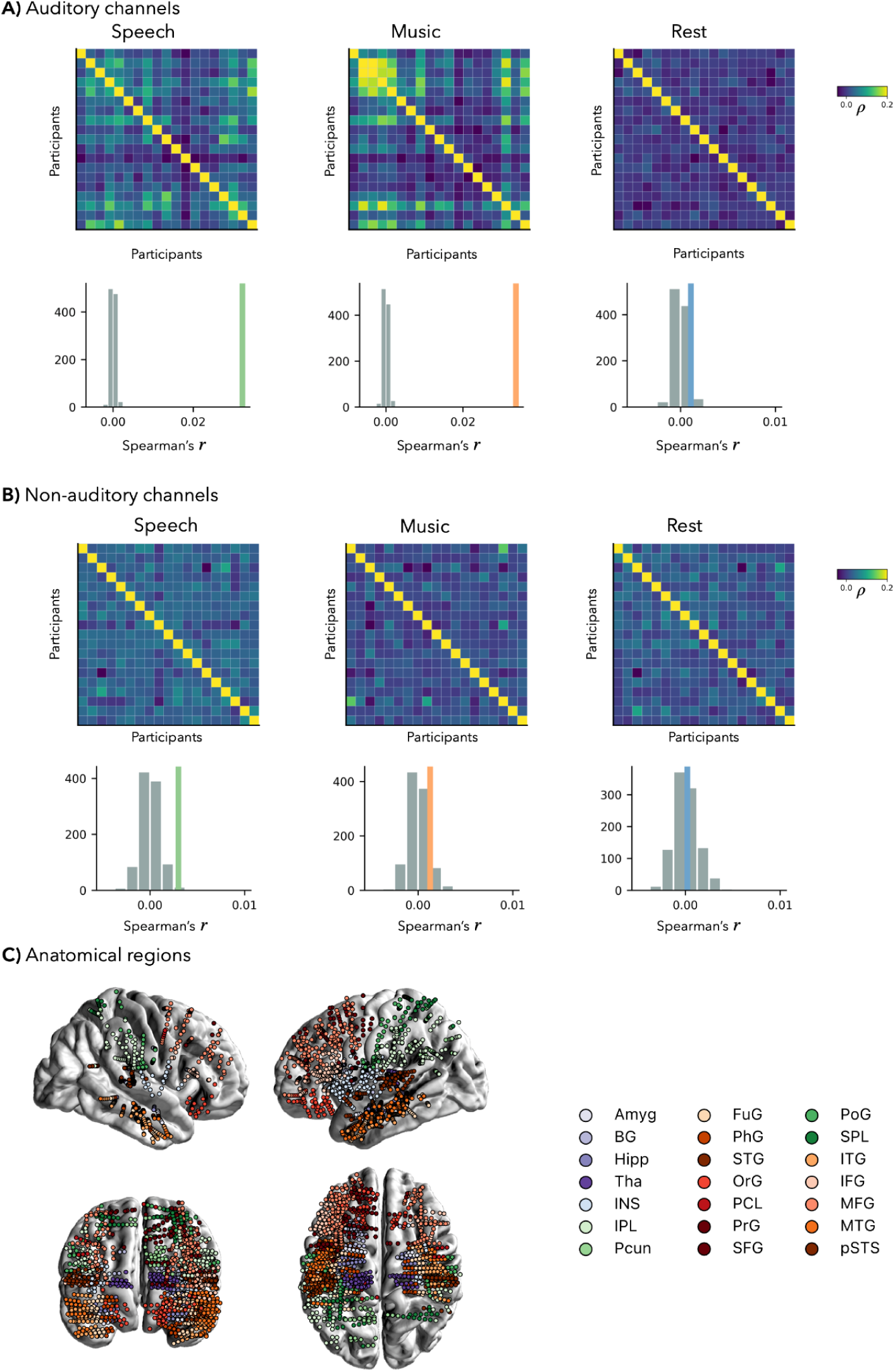
Inter-subject correlation of the APs across conditions. Mean of the inter-subject correlation (coloured line), compared to a null model distribution (see Methods) for **(A)** auditory only electrodes (H) and **(B)** non-auditory only electrodes (∼H). **(C)** Anatomical regions defined according to the Brainnetome atlas (57). Abbreviations according to the Brainnetome atlas.

**Supplementary Figure S2.**
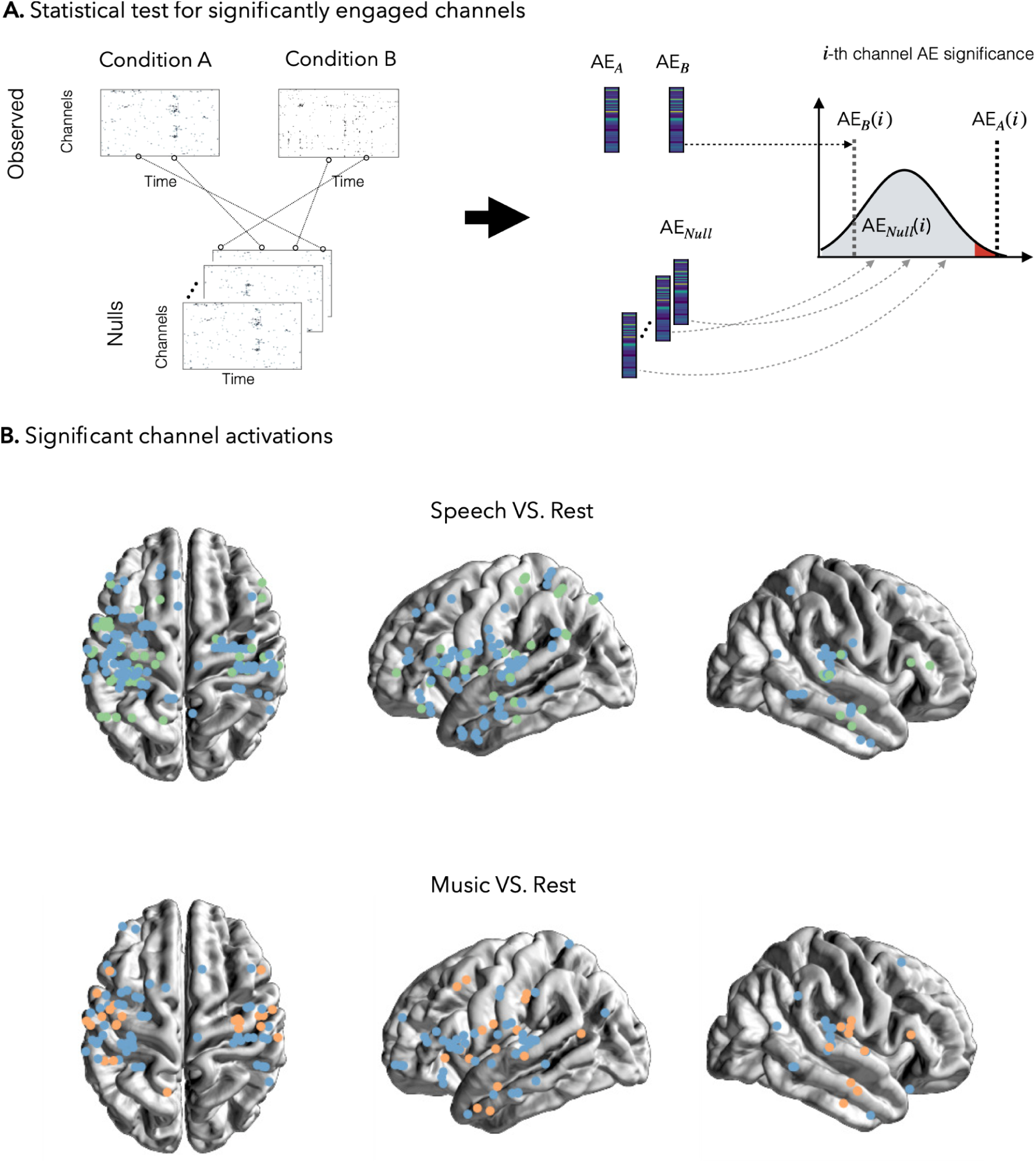
**(A)** The statistical test for comparing AE across conditions consists of 1) defining null binary sequences from the random combination of binary signals observed in conditions A and B (among speech, music, and rest); 2) extracting the AE for each null data and defining the (hypergeometric) null-distribution for each channel *i*; 3) compare the AE(*i*) observed in conditions A and B to the null-distribution of AE_*Null*_(*i*). Significantly engaged channels display AE(*i*) falling in the right tail of the null distribution (p<0.05, Bonferroni corrected for the number of channels). **(B)** Significantly engaged channels across all pairs of conditions (speech in green, music in orange, rest in blue) and across all participants.

**Supplementary Figure S3.**
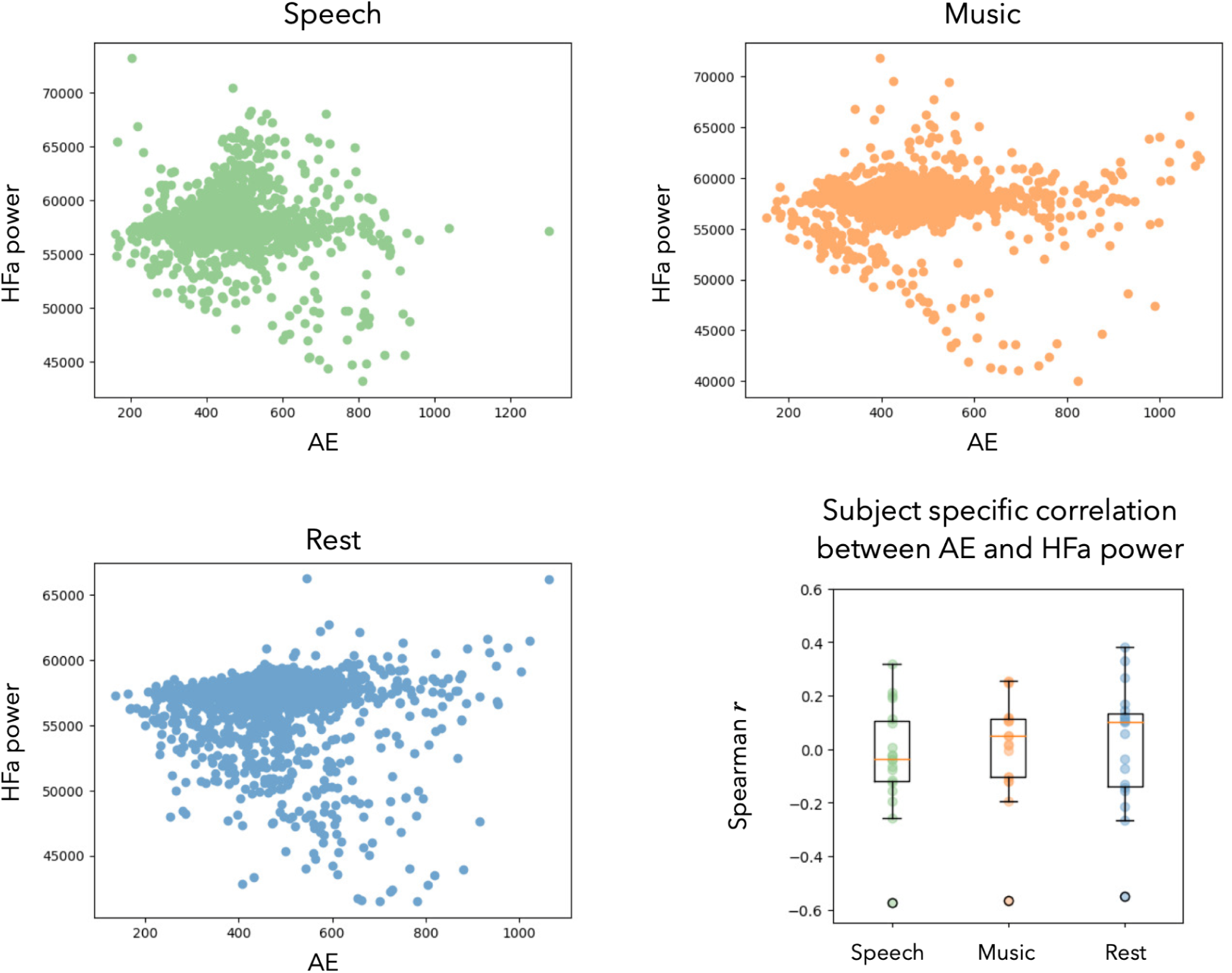
The three scatterplots show–for all channels present in our data –the HFa power, measured as the sum of the original filtered signals (not z-scored), and the AE, computed as the sum of the binarized (z-scored) signals. At the level of aggregate population these scatterplots show poor correlation values in all conditions (Speech, Spearman r= -0.00039, p=0.98; Music, Spearman r= -0.0558, p<0.05; Rest, Spearman r= -0.0765, p<0.05). When computed for each participant separately (bottom right panel), some correlation values between AE and HFa power were higher in absolute value, but the distribution of correlation values was not significantly deviating from zero (one sample t-test p>0.05).

### Appendix A: Neuronal avalanches association to cortical timescales

In this section we show that the AP and AE measures obtained from the binarized activity can be associated with signals autocorrelation features, and interpreted as a proxy for the dynamics and topography of neuronal timescales.

In general, a characteristic timescale can be defined for a signal by studying the decay time of its auto-correlation function. Recent studies demonstrate that the brain’s cortical timescales (e.g., measured by autocorrelation decay time) follow a smooth gradient spanning from posterior sensorimotor regions to anterior trans-modal hubs (60). Heterogeneous structural features such as brain microstructure, connectivity, cortical thickness, and gene expression, vary along the same axis (61,62), which points to a deep link between structure and function (63,64). This hierarchical structure is preserved during the course of mammalian evolution, despite the increase of brain volume (62); it is characteristic of individual brains, and alterations in its organization lead to mental and neurological disease (65). However, differently than structural features, brain function is highly dynamic and the topography of timescales can vary over time both in rest (66), and task (67). For instance, visual stimuli presentation induces a decrease in the amplitude of the alpha band (8-12Hz), while mental calculation or working memory tasks increase the activity in this same frequency band (7,68).

While several measures for the signals timescale have been proposed (64,66), here for illustrative purposes we evaluated the timescales as the area τ before the first minimum; Fig.S4.A, left). Even in the same recording channel, the decay of the auto-correlation can change when observed in different time windows (Fig.S4.A, right). To compare the timescales results to the neuronal avalanches results obtained in Fig.1, we first studied the time evolution of the timescale τ_*i*_(t) for each channel *i* (Fig.S4.B). Averaging across all time windows, we defined the Temporal Engagement (TE) showing that electrophysiological fluctuations at different channels are associated with different timescales on average (Fig.S4.B, right). Averaging across channels, we define the Temporal Profile (TP) which shows that the system’s timescales can be shorter or longer depending on the observation window (Fig.S4.B, bottom). Notably, both the TE and the TP correlate with the activity engagement (AE) and activity profile (AP) (averaged in time windows to allow comparison), which were previously defined from the binarized activity (Fig.S4.C). This shows firstly that the fluctuations of the system’s time scales are captured by the avalanche approach, secondly that the spatial characterization of the timescales, e.g., in which brain regions are short/long timescales observed on average, depends on how many times the avalanches were passing from each brain region. Importantly, using the decay of the autocorrelation as a measure of dynamic timescales we obtained similar inter-subject correlation results as in the binary approach (Fig.S4.D). More specifically we have a significant inter-subject correlation during speech listening (p < 0.001) and during music listening (p = 0.02) and not significant during resting state (p = 0.877). The significance of the inter-subject correlation between TPs is assessed by shifting 1000 iterations as for the APs.

Since we show that both the time profile of binary activations (Activity Profile AP) and the spatial topography of activations (Activity Engagement AE) are highly correlated with related temporal (Time Profile TP) and spatial (Time Engagement TE) metrics obtained from spectral features, the presented metrics could be used in the future to analyze timescale dynamics at a fine-grained temporal resolution and low computational cost. Overall, the observation of increased neuronal avalanche activity in association with slower neuronal timescales provide evidence for neuronal avalanches being involved in slow top-down processing of information.

**Figure S4.**
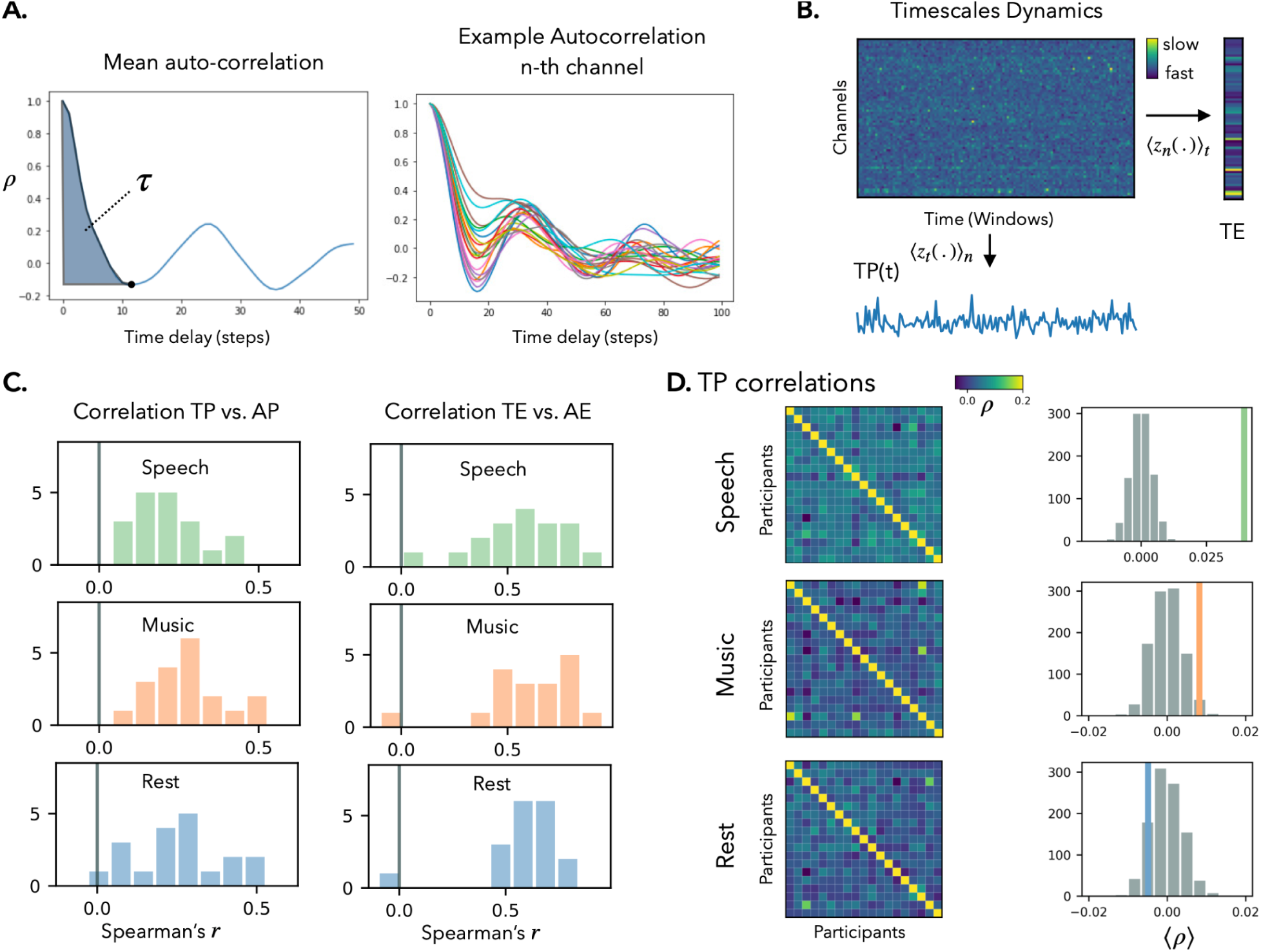
**A, left)** A timescale τ for a brain signal was defined as the area under the autocorrelation plot, between 0 and the first minimum. For each channel, this plot can be obtained and τ can be computed in different time windows. **A, right)** In a single channel, the autocorrelation decay time can vary over time (each color represents different time windows). **B)** For each channel the timescales are computed as explained in (A) in windows of 150 time steps (3 seconds) and their evolution in time is investigated using a sliding window approach. The temporal evolution of the timescales is plotted in channels x time windows fashion. The Temporal Profile (TP) and Temporal Engagement (TE), computed by summing the timescales dynamics matrix across channels and time, respectively, are depicted below and on the right side of the matrix. **(C)** The distribution of the correlation values between the APs and TPs (left side), and between the AEs and TEs (right side) in each condition. **D, left)** Inter-subject correlation matrices computed on the TPs. **D, right)** the comparison with the null model, computed randomly by time-shifting the TPs is shown and a statistic is computed.

**Table S1.**
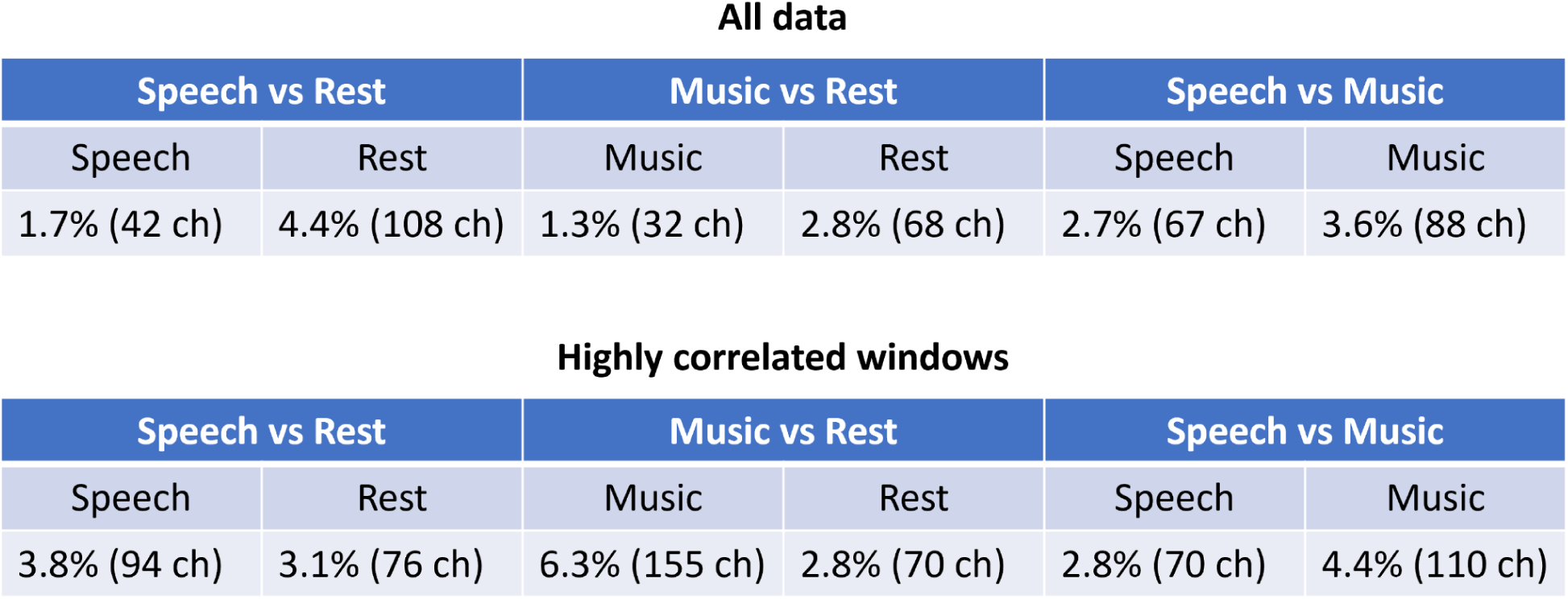
Percentage of high activity engagement (high-AE) channels comparing all conditions (speech, music, and rest) in pairs. **Top table)** the percentages are computed over all the data points. **Bottom table)** the percentages of high-AE for speech and music are computed considering only the time windows presenting intersubject correlation higher than the maximum observed during resting state (red dots in Fig.2.B).

All data
| Speech vs Rest |  | Music vs Rest |  | Speech vs Music |  |
| --- | --- | --- | --- | --- | --- |
| Speech | Rest | Music | Rest | Speech | Music |
| 1.7% (42 ch) | 4.4% (108 ch) | 1.3% (32 ch) | 2.8% (68 ch) | 2.7% (67 ch) | 3.6% (88 ch) |

**Table S1.** *Percentage of high activity engagement (high-AE) channels comparing all conditions (speech, music, and rest) in pairs.
| Speech vs Rest |  | Music vs Rest |  | Speech vs Music |  |
| --- | --- | --- | --- | --- | --- |
| Speech | Rest | Music | Rest | Speech | Music |
| 3.8% (94 ch) | 3.1% (76 ch) | 6.3% (155 ch) | 2.8% (70 ch) | 2.8% (70 ch) | 4.4% (110 ch) |

## Bibliography

1. Bressler SL, Menon V. Large-scale brain networks in cognition: emerging methods and principles. Trends Cogn Sci (Regul Ed). 2010 Jun;14(6):277–90.

2. Park H-J, Friston K. Structural and functional brain networks: from connections to cognition. Science. 2013 Nov 1;342(6158):1238411.

3. te Rietmolen N, Mercier M, Trébuchon A, Morillon B, Schön D. Speech and music recruit frequency-specific distributed and overlapping cortical networks. BioRxiv. 2022 Oct 9;

4. Buzsáki G, Watson BO. Brain rhythms and neural syntax: implications for efficient coding of cognitive content and neuropsychiatric disease. Dialogues Clin Neurosci. 2012 Dec;14(4):345–67.

5. Bullmore E, Sporns O. Complex brain networks: Graph theoretical analysis of structural and functional systems. Nat Rev Neurosci. 2009 Mar;10(3):186–98.

6. Beggs JM, Plenz D. Neuronal avalanches in neocortical circuits. J Neurosci. 2003 Dec 3;23(35):11167–77.

7. Palva JM, Zhigalov A, Hirvonen J, Korhonen O, Linkenkaer-Hansen K, Palva S. Neuronal long-range temporal correlations and avalanche dynamics are correlated with behavioral scaling laws. Proc Natl Acad Sci USA. 2013 Feb 26;110(9):3585–90.

8. Priesemann V, Valderrama M, Wibral M, Le Van Quyen M. Neuronal avalanches differ from wakefulness to deep sleep--evidence from intracranial depth recordings in humans. PLoS Comput Biol. 2013 Mar 21;9(3):e1002985.

9. Tagliazucchi E, Balenzuela P, Fraiman D, Chialvo DR. Criticality in large-scale brain FMRI dynamics unveiled by a novel point process analysis. Front Physiol. 2012 Feb 8;3:15.

10. Shriki O, Alstott J, Carver F, Holroyd T, Henson RNA, Smith ML, et al. Neuronal avalanches in the resting MEG of the human brain. J Neurosci. 2013 Apr 17;33(16):7079–90.

11. Bak P, Tang C, Wiesenfeld K. Self-organized criticality. Phys Rev A. 1988 Jul 1;38(1):364–74.

12. di Santo S, Villegas P, Burioni R, Muñoz MA. Landau-Ginzburg theory of cortex dynamics: Scale-free avalanches emerge at the edge of synchronization. Proc Natl Acad Sci USA. 2018 Feb 13;115(7):E1356–65.

13. Buendía V, Villegas P, Burioni R, Muñoz MA. Hybrid-type synchronization transitions: Where incipient oscillations, scale-free avalanches, and bistability live together. Phys Rev Research. 2021 Jun 21;3(2):023224.

14. Plenz D, Ribeiro TL, Miller SR, Kells PA, Vakili A, Capek EL. Self-Organized Criticality in the Brain. Front Phys. 2021 Jul 7;9.

15. Shew WL, Yang H, Yu S, Roy R, Plenz D. Information capacity and transmission are maximized in balanced cortical networks with neuronal avalanches. J Neurosci. 2011 Jan 5;31(1):55–63.

16. Bansal K, Garcia JO, Lauharatanahirun N, Muldoon SF, Sajda P, Vettel JM. Scale-specific dynamics of high-amplitude bursts in EEG capture behaviorally meaningful variability. Neuroimage. 2021 Nov 1;241:118425.

17. Pajevic S, Plenz D. Efficient network reconstruction from dynamical cascades identifies small-world topology of neuronal avalanches. PLoS Comput Biol. 2009 Jan 30;5(1):e1000271.

18. Zhou X, Ma N, Song B, Wu Z, Liu G, Liu L, et al. Optimal Organization of Functional Connectivity Networks for Segregation and Integration With Large-Scale Critical Dynamics in Human Brains. Front Comput Neurosci. 2021 Mar 31;15:641335.

19. Beggs JM, Timme N. Being critical of criticality in the brain. Front Physiol. 2012 Jun 7;3:163.

20. Turkheimer FE, Rosas FE, Dipasquale O, Martins D, Fagerholm ED, Expert P, et al. A complex systems perspective on neuroimaging studies of behavior and its disorders. Neuroscientist. 2022 Aug;28(4):382–99.

21. Seshadri S, Klaus A, Winkowski DE, Kanold PO, Plenz D. Altered avalanche dynamics in a developmental NMDAR hypofunction model of cognitive impairment. Transl Psychiatry. 2018 Jan 10;8(1):3.

22. Duma GM, Danieli A, Mento G, Vitale V, Opipari RS, Jirsa V, et al. Altered spreading of neuronal avalanches in temporal lobe epilepsy relates to cognitive performance: A resting-state hdEEG study. Epilepsia. 2023 May;64(5):1278–88.

23. Rucco R, Bernardo P, Lardone A, Baselice F, Pesoli M, Polverino A, et al. Neuronal Avalanches to Study the Coordination of Large-Scale Brain Activity: Application to Rett Syndrome. Front Psychol. 2020 Oct 27;11:550749.

24. Polverino A, Troisi Lopez E, Minino R, Liparoti M, Romano A, Trojsi F, et al. Flexibility of fast brain dynamics and disease severity in amyotrophic lateral sclerosis. Neurology. 2022 Nov 22;99(21):e2395–405.

25. Sorrentino P, Rucco R, Baselice F, De Micco R, Tessitore A, Hillebrand A, et al. Flexible brain dynamics underpins complex behaviours as observed in Parkinson’s disease. Sci Rep. 2021 Feb 18;11(1):4051.

26. Scarpetta S, Morrisi N, Mutti C, Azzi N, Trippi I, Ciliento R, et al. Criticality of neuronal avalanches in human sleep and their relationship with sleep macro- and micro-architecture. iScience. 2023 Oct 20;26(10):107840.

27. Finn ES. Is it time to put rest to rest? Trends Cogn Sci (Regul Ed). 2021 Dec;25(12):1021–32.

28. Hasson U, Simmons JP, Todorov A. Believe it or not: on the possibility of suspending belief. Psychol Sci. 2005 Jul;16(7):566–71.

29. Sonkusare S, Breakspear M, Guo C. Naturalistic stimuli in neuroscience: critically acclaimed. Trends Cogn Sci (Regul Ed). 2019 Aug;23(8):699–714.

30. Hasson U, Malach R, Heeger DJ. Reliability of cortical activity during natural stimulation. Trends Cogn Sci (Regul Ed). 2010 Jan;14(1):40–8.

31. Dini H, Simonetti A, Bruni LE. Exploring the Neural Processes behind Narrative Engagement: An EEG Study. eNeuro. 2023 Jul 17;10(7).

32. Hamilton LS, Huth AG. The revolution will not be controlled: natural stimuli in speech neuroscience. Lang Cogn Neurosci. 2020;35(5):573–82.

33. Corsi M-C, Sorrentino P, Schwartz D, George N, Hugueville L, Kahn AE, et al. Measuring brain critical dynamics to inform Brain-Computer Interfaces. BioRxiv. 2022 Jun 17;

34. Ray S, Maunsell JHR. Different origins of gamma rhythm and high-gamma activity in macaque visual cortex. PLoS Biol. 2011 Apr 12;9(4):e1000610.

35. Martin S, Brunner P, Iturrate I, Millán JDR, Schalk G, Knight RT, et al. Word pair classification during imagined speech using direct brain recordings. Sci Rep. 2016 May 11;6:25803.

36. Mercier MR, Dubarry A-S, Tadel F, Avanzini P, Axmacher N, Cellier D, et al. Advances in human intracranial electroencephalography research, guidelines and good practices. Neuroimage. 2022 Oct 15;260:119438.

37. Lannelongue L, Inouye M. Carbon footprint estimation for computational research. Nat Rev Methods Primers. 2023 Feb 16;3(1):9.

38. Fedorenko E, Thompson-Schill SL. Reworking the language network. Trends Cogn Sci (Regul Ed). 2014 Mar;18(3):120–6.

39. Fontenele AJ, Sooter JS, Norman VK, Gautam SH, Shew WL. Low dimensional criticality embedded in high dimensional awake brain dynamics. BioRxiv. 2023 Jul 24;

40. Sorrentino P, Seguin C, Rucco R, Liparoti M, Troisi Lopez E, Bonavita S, et al. The structural connectome constrains fast brain dynamics. eLife. 2021 Jul 9;10.

41. Buzsáki G, Moser EI. Memory, navigation and theta rhythm in the hippocampal-entorhinal system. Nat Neurosci. 2013 Feb;16(2):130–8.

42. Buzsáki G. Rhythms of the Brain. Oxford University Press; 2006.

43. Fries P. Rhythms for Cognition: Communication through Coherence. Neuron. 2015 Oct 7;88(1):220–35.

44. van Ede F, Quinn AJ, Woolrich MW, Nobre AC. Neural Oscillations: Sustained Rhythms or Transient Burst-Events? Trends Neurosci. 2018 Jul;41(7):415–7.

45. Heims SJ. The Cybernetics Group. The MIT Press; 1991.

46. Timme NM, Lapish C. A tutorial for information theory in neuroscience. eNeuro. 2018 Sep 11;5(3).

47. Bernard C. Brain’s best kept secret: degeneracy. eNeuro. 2023 Nov 14;10(11).

48. Edelman GM, Gally JA. Degeneracy and complexity in biological systems. Proc Natl Acad Sci USA. 2001 Nov 20;98(24):13763–8.

49. Price CJ. The anatomy of language: contributions from functional neuroimaging. J Anat. 2000 Oct;197 Pt 3(Pt 3):335–59.

50. Warren J. How does the brain process music? Clin Med. 2008 Feb;8(1):32–6.

51. Peretz I, Vuvan D, Lagrois M-É, Armony JL. Neural overlap in processing music and speech. Philos Trans R Soc Lond B Biol Sci. 2015 Mar 19;370(1664):20140090.

52. Lachaux J-P, Axmacher N, Mormann F, Halgren E, Crone NE. High-frequency neural activity and human cognition: past, present and possible future of intracranial EEG research. Prog Neurobiol. 2012 Sep;98(3):279–301.

53. Taylor TJ, Hartley C, Simon PL, Kiss IZ, Berthouze L. Identification of Criticality in Neuronal Avalanches: I. A Theoretical Investigation of the Non-driven Case. J Math Neurosci. 2013 Apr 23;3(1):5.

54. Cocchi L, Gollo LL, Zalesky A, Breakspear M. Criticality in the brain: A synthesis of neurobiology, models and cognition. Prog Neurobiol. 2017 Nov;158:132–52.

55. Mercier MR, Bickel S, Megevand P, Groppe DM, Schroeder CE, Mehta AD, et al. Evaluation of cortical local field potential diffusion in stereotactic electro-encephalography recordings: A glimpse on white matter signal. Neuroimage. 2017 Feb 15;147:219–32.

56. Groppe DM, Bickel S, Dykstra AR, Wang X, Mégevand P, Mercier MR, et al. iELVis: An open source MATLAB toolbox for localizing and visualizing human intracranial electrode data. J Neurosci Methods. 2017 Apr 1;281:40–8.

57. Fan L, Li H, Zhuo J, Zhang Y, Wang J, Chen L, et al. The human brainnetome atlas: A new brain atlas based on connectional architecture. Cereb Cortex. 2016 Aug;26(8):3508–26.

58. Ossandón T, Vidal JR, Ciumas C, Jerbi K, Hamamé CM, Dalal SS, et al. Efficient “pop-out” visual search elicits sustained broadband γ activity in the dorsal attention network. J Neurosci. 2012 Mar 7;32(10):3414–21.

59. Vidal JR, Freyermuth S, Jerbi K, Hamamé CM, Ossandon T, Bertrand O, et al. Long-distance amplitude correlations in the high γ band reveal segregation and integration within the reading network. J Neurosci. 2012 May 9;32(19):6421–34.

60. Murray JD, Bernacchia A, Freedman DJ, Romo R, Wallis JD, Cai X, et al. A hierarchy of intrinsic timescales across primate cortex. Nat Neurosci. 2014 Dec;17(12):1661–3.

61. Gao R, van den Brink RL, Pfeffer T, Voytek B. Neuronal timescales are functionally dynamic and shaped by cortical microarchitecture. eLife. 2020 Nov 23;9.

62. Fulcher BD, Murray JD, Zerbi V, Wang X-J. Multimodal gradients across mouse cortex. Proc Natl Acad Sci USA. 2019 Mar 5;116(10):4689–95.

63. Arbabshirani MR, Damaraju E, Phlypo R, Plis S, Allen E, Ma S, et al. Impact of autocorrelation on functional connectivity. Neuroimage. 2014 Nov 15;102 Pt 2(0 2):294–308.

64. Müller PM, Meisel C. Spatial and temporal correlations in human cortex are inherently linked and predicted by functional hierarchy, vigilance state as well as antiepileptic drug load. PLoS Comput Biol. 2023 Mar 3;19(3):e1010919.

65. Northoff G, Sandsten KE, Nordgaard J, Kjaer TW, Parnas J. The self and its prolonged intrinsic neural timescale in schizophrenia. Schizophr Bull. 2021 Jan 23;47(1):170–9.

66. Sorrentino P, Rabuffo G, Baselice F, Troisi Lopez E, Liparoti M, Quarantelli M, et al. Dynamical interactions reconfigure the gradient of cortical timescales. Netw Neurosci. 2023 Jan 1;7(1):73–85.

67. Golesorkhi M, Gomez-Pilar J, Çatal Y, Tumati S, Yagoub MCE, Stamatakis EA, et al. From temporal to spatial topography: hierarchy of neural dynamics in higher- and lower-order networks shapes their complexity. Cereb Cortex. 2022 Dec 8;32(24):5637–53.

68. Palva S, Palva JM. New vistas for alpha-frequency band oscillations. Trends Neurosci. 2007 Apr;30(4):150–8.

